# Characterization of *Streptococcus pneumoniae* phage-like element SpnCI reveals an enhanced virulent phenotype in the acute invertebrate infection model *Galleria mellonella*

**DOI:** 10.1101/670141

**Authors:** Kimberly McCullor, Maliha Rahman, Catherine King, W. Michael McShan

## Abstract

Phage-like elements are found in a multitude of streptococcal species, including pneumococcal strain Hungary19A-6 (SpnCI). The aim of our research was to investigate the role of phage-like element SpnCI in enhanced virulence and phenotypic modulation within *Streptococcus pneumoniae*. SpnCI was found to significantly enhance virulence within the invertebrate infection model *Galleria mellonella*. Infections with SpnCI led to a lower mean health score (1.6) and survival percentage (20%) compared to SpnCI null TIGR4 infections (3.85 mean health score and 50% survival). SpnCI remained integrated throughout growth, conferring greater sensitivity to UV irradiation. Change in transcriptional patterns occurred, including downregulation of operons involved with cell surface modelling in the SpnCI containing strain of TIGR4. Kanamycin-tagged SpnCI strain in Hungary19A-6 was inducible and isolated from lysate along with both annotated prophages. No phages were identified by PCR nor electron microscopy (EM) following induction of TIGR4 SpnCIΔ*strA* suggesting helper-phage dependence for dissemination. EM of lysate showed typical siphoviridae morphology with an average capsid size of 60 nm. Two of sixty capsids were found to be smaller, suggesting SpnCI disseminates using a similar mechanism described for *Staphylococcus aureus* phage-like element SaPI. SpnCI from lysate infected capsule null strain T4R but was incapable of infecting the encapsulated TIGR4 strain suggesting that capsule impedes phage infection. Our work demonstrates that SpnCI can modulate virulence, UV susceptibility, alter transcriptional patterns, and furthermore, can disseminate via infection within pneumococcus. Further research is necessary to elucidate how SpnCI modulates virulence and what genes are responsible for the enhanced virulence phenotype.

**Importance:** Although vaccines have limited the scope of pneumococcal infections, *Streptococcus pneumoniae* still remains an important human pathogen. Understanding novel elements, such as SpnCI, that enhance virulence can lead to the development of more targeted therapeutic and diagnostic tools within the clinical realm.

## Introduction

*Streptococcus pneumoniae*, the major causative agent of pediatric otitis media and adult pneumonia, still remains an important human pathogen even with the advent of vaccines. *S. pneumoniae* is responsible for over 400,000 hospitalizations annually in the United States (1), with 30% of infections being resistant to one or more commonly used antibiotics. Pneumococcal infections are estimated to place a $96 million per year burden on the healthcare system in the United States alone (2, 3). Our group was the first to identify streptococcal phage-like chromosomal islands (PLCI). Previously we have demonstrated that in *Streptococcus pyogenes*, phage-like element SpyCIM1 (***<u>S</u>**treptococcus **<u>py</u>**ogenes* **<u>C</u>**hromosomal **<u>I</u>**sland **<u>M1</u>** strain) led to a growth dependent mutator phenotype (due to the disruption of DNA mismatch repair (MMR) operon through SpyCIM1 site-specific integration during late logarithmic to stationary phase of growth) (4, 5). We have also shown that SpyCIM1 could confer global transcriptional changes leading to up regulation of known virulence genes such as *hasB* and *emm* (6) and was found to correspond to an enhanced virulent phenotype within the invertebrate acute infection model *Galleria mellonella* (unpublished). *S. anginosus* phage-like element SanCI was found to enhance virulence in the *G. mellonella* infection model as well (unpublished) suggesting that phage-like chromosomal islands may serve as virulence factors. We have found through *in silico* analysis the presence of phage-like chromosomal islands in a multitude of streptococcal species with the majority integrating within the MMR operon like SpyCIM1 or within other sites involved with DNA repair mechanisms such as nucleotide excision repair (NER) (the integration site for *S. pneumoniae* phage-like chromosomal island, SpnCI) (7). *S. pneumoniae* multi-drug resistant strain Hungary 19A-6 (8), was found to contain this particular phage-like chromosomal island, SpnCI. The 19A serotype is correlated with a higher prevalence of invasive disease and increased antimicrobial resistance especially noted after the introduction of the conjugate pneumococcal vaccine (9-17). Some reports describe 19A emergence within countries where PCV7 was not introduced (18), suggesting that vaccine selective pressure is not the only driver to 19A emergence. Here we demonstrate that SpnCI enhances virulence using the invertebrate acute infection model *Galleria mellonella* (19), confers UV susceptibility due to maintained integration (suggesting loss of NER), and its capabilities for altering transcriptional patterns. We also describe the dissemination of SpnCI beyond transformation. As more appreciation of the role phage-like chromosomal islands play in gene expression and evolution in Gram-positive and Gram-negative organisms (20-23), a better understanding of SpnCI and its role in virulence and potential for conferring beneficial mutations provides a novel dynamic in *Streptococcus pneumoniae* infections.

## Results

### SpnCI: A Phage-like Chromosomal Island of *Streptococcus pneumoniae*

Genome database mining for streptococcal PLCI discovered such an element in *S. pneumoniae strain* Hungary19A-6 (Genbank accession NC_010380) that is integrated into the overlapping promoters of *corA* and *uvrA*, which encodes the A subunit of the UvrABC excinuclease required for NER. Following our previous convention, the element was named SpnCI (**<u>S</u>**treptococcus **<u>p</u>**neumoniae **<u>C</u>**hromosomal **<u>I</u>**sland). Figure 1A shows the integrated SpnCI as found in the bacterial genome; the genes for integrase (*int*) and *strA* are indicated. Irradiation of the bacterium with ultraviolet light (UV) causes the SpnCI to excise from the chromosome, restoring the native configuration of the corA-uvrA promoter region (*attB*) as well as the junction of the circular form of SpnCI (*attP*, Figure 1B). Thus, SpnCI has the potential to act as a molecular switch for genes *corA* and *uvrA* depending upon its integration state.

**Figure 1.**
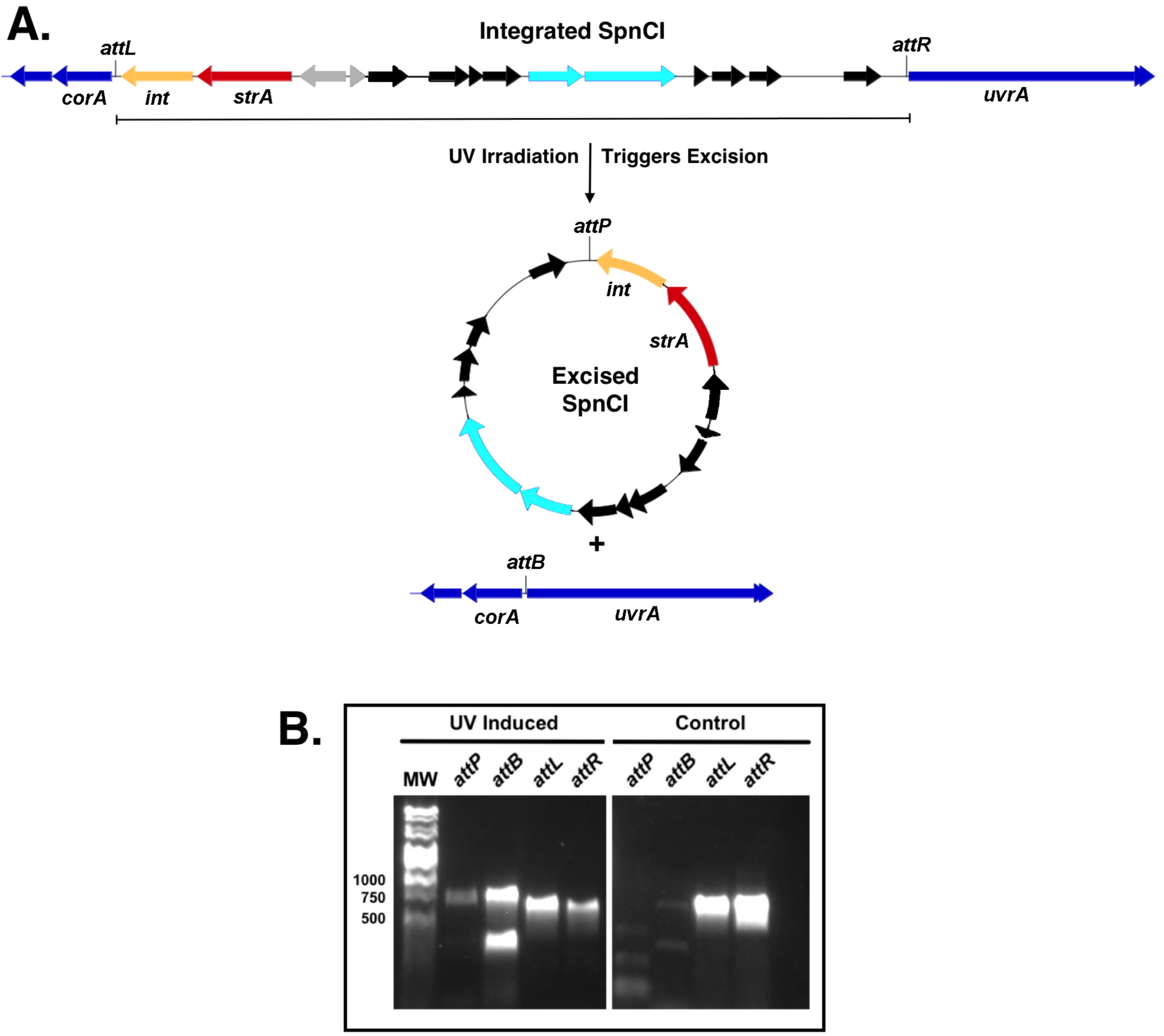
SpnCI is able to excise and integrate into the *S. pneumoniae* genome. **A.** *S. anginosus* phage-like SanCI shares a similar fusion peptide *strA* (red) that consists of a helix-turn-helix motif and predicted ATPase region. Interestingly, this peptide is not found in *S. pyogenes* phage-like element SpyCI. Other predicted SpnCI genes include those involved in regulating lysogeny (grey) and DNA replication (light blue) but the majority of the genes contained are of unknown function (black). **B.** UV treatment induces excision of phage-like element SpnCI from the shared promoter site of *corA* (heavy metal transporter) and *uvrA* (nucleotide excision repair) (see *attB* site) within the *S. pneumoniae* genome (dark blue). Both *attP* and *attB* are amplified post UV treatment compared to the control where the integrated state (*attL* and *attR*) is predominantly amplified. This confirms SpnCI contains a functioning integrase (yellow) and excisionase (gene identity unknown).

### Introducing SpnCI into the TIGR4 genetic background

To create a pair of isogenic strains that differed only by the presence or absence of SpnCI, the PLCI from *S. pneumoniae* strain Hungary 19A-6 was first tagged with a selectable marker by replacing gene *strA* with a kanamycin resistance cassette via homologous recombination of the flanking region (Figure 2A). The tagged SpnCI was then moved into capsule null strain T4R, a derivative of *S. pnuemoniae* TIGR4. The capsule was rescued in the resulting SpnCI+ strain by transforming T4R SpnCI Δ*strA* with wild type *cps4A* in order to generate TIGR4 SpnCIΔ*strA* (Figure 2B).

**Figure 2.**
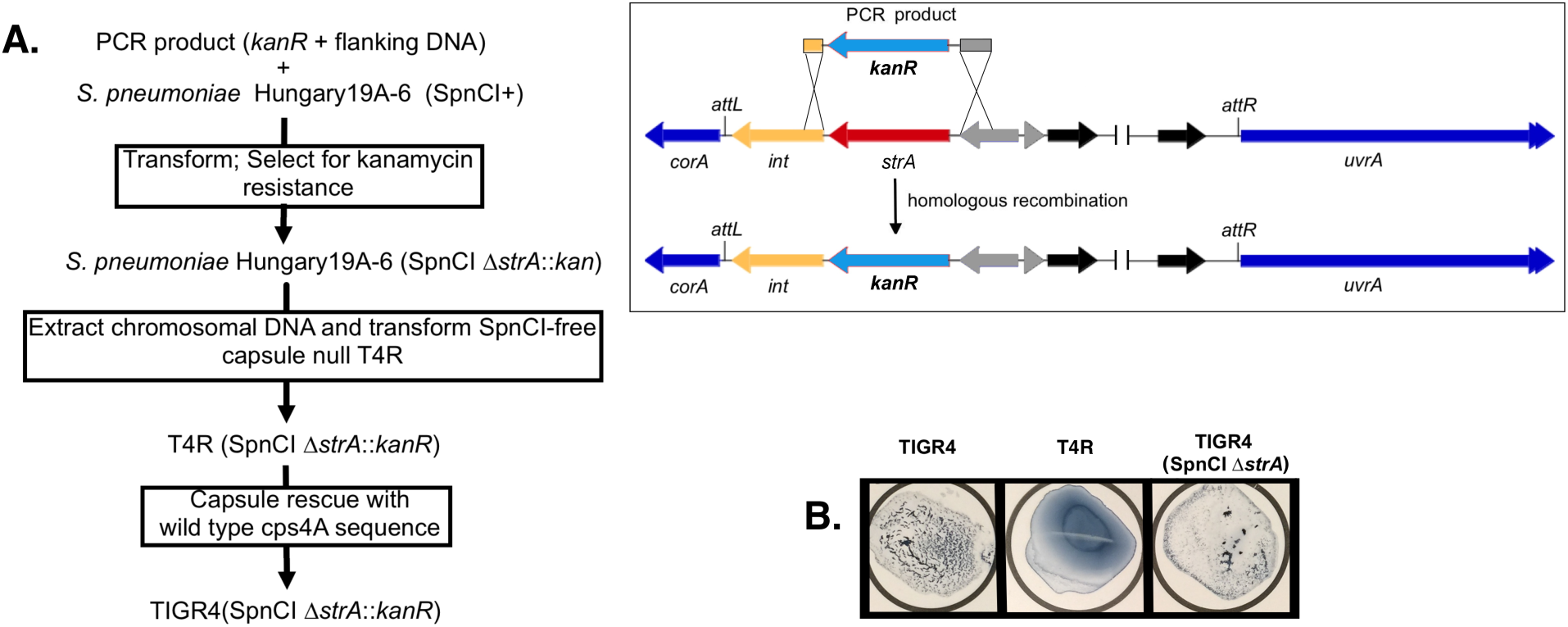
Strategy for introducing SpnCI into the TIGR4 genetic background. **A.** Strategy employed to move SpnCI into strain TIGR4. SpnCI was first tagged with kanamycin resistance (*kanR*) and moved into capsule null TIGR4 background strain T4R. Capsule was then rescued by transformation with wildtype *cps4A* to rescue the capsule phenotype and to generate the TIGR4 SpnCIΔ*strA* strain. **B.** Restored capsule expression was confirmed by agglutination test and by sequencing the rescued wild type *cps4A* loci.

### SpnCI integration and impact on UV susceptibility

To investigate the abundance of integrated or excised states of SpnCI, Hungary19A-6 DNA was collected throughout the growth cycle (A_600_=0.1 through 0.8). Studies demonstrated a mixed culture of integrated and excised SpnCI states throughout the early log to stationary phase of growth by PCR. Further quantitative analysis via qPCR demonstrated that the majority of SpnCI remained integrated at each time point tested as the fold change remained constant (around 1) throughout the bacterial growth cycle for *attL* and *attR* when compared to the initial 15-minute time point (Figure 3). Sites *attP* and *attB* failed to amplify or their Ct values were extremely high indicating a small population existed in an excised state. As integration interrupts the shared promoter site for divergent genes *corA* (heavy metal transporter) and *uvrA* (nucleotide excision repair component), we next investigated UV susceptibility amongst the isogenic strains of TIGR4 with and without SpnCI using the TIGR4 SpnCIΔ*strA* generated strain. The SpnCI containing strain was observed to be more susceptible to UV compared to wild type. This is especially evident at late log time point (O.D. _A600_= 0.6). The addition of SpnCI alone made the TIGR4 genetic background more similar to wildtype SpnCI containing strain Hungary19A-6 which is markedly more susceptible to UV damage (Figure 4).

**Figure 3.**
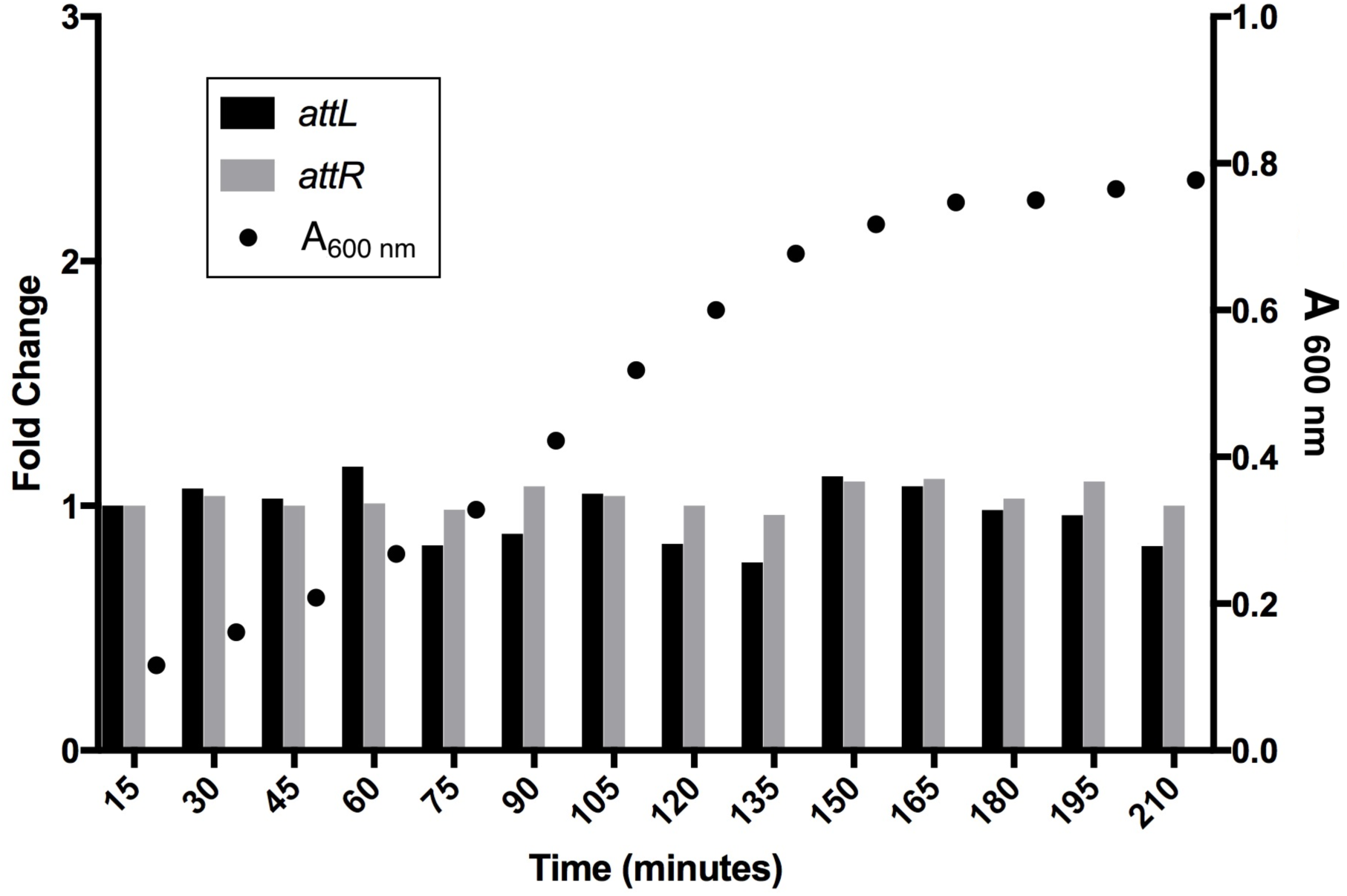
SpnCI Excision is not dependent on growth and remains predominantly integrated. Fold change is in comparison to the initial 15-minute time point at early logarithmic growth for *attL* (black) and *attR* (grey). Nominal differences in fold change was observed throughout growth indicating that SpnCI remains integrated throughout logarithmic and stationary phase (optical density of bacterial cultures shown on left y-axis; circles). Little to no amplification was observed for excised specific qPCR primers *attP* and *attB* (data not shown).

**Figure 4.**
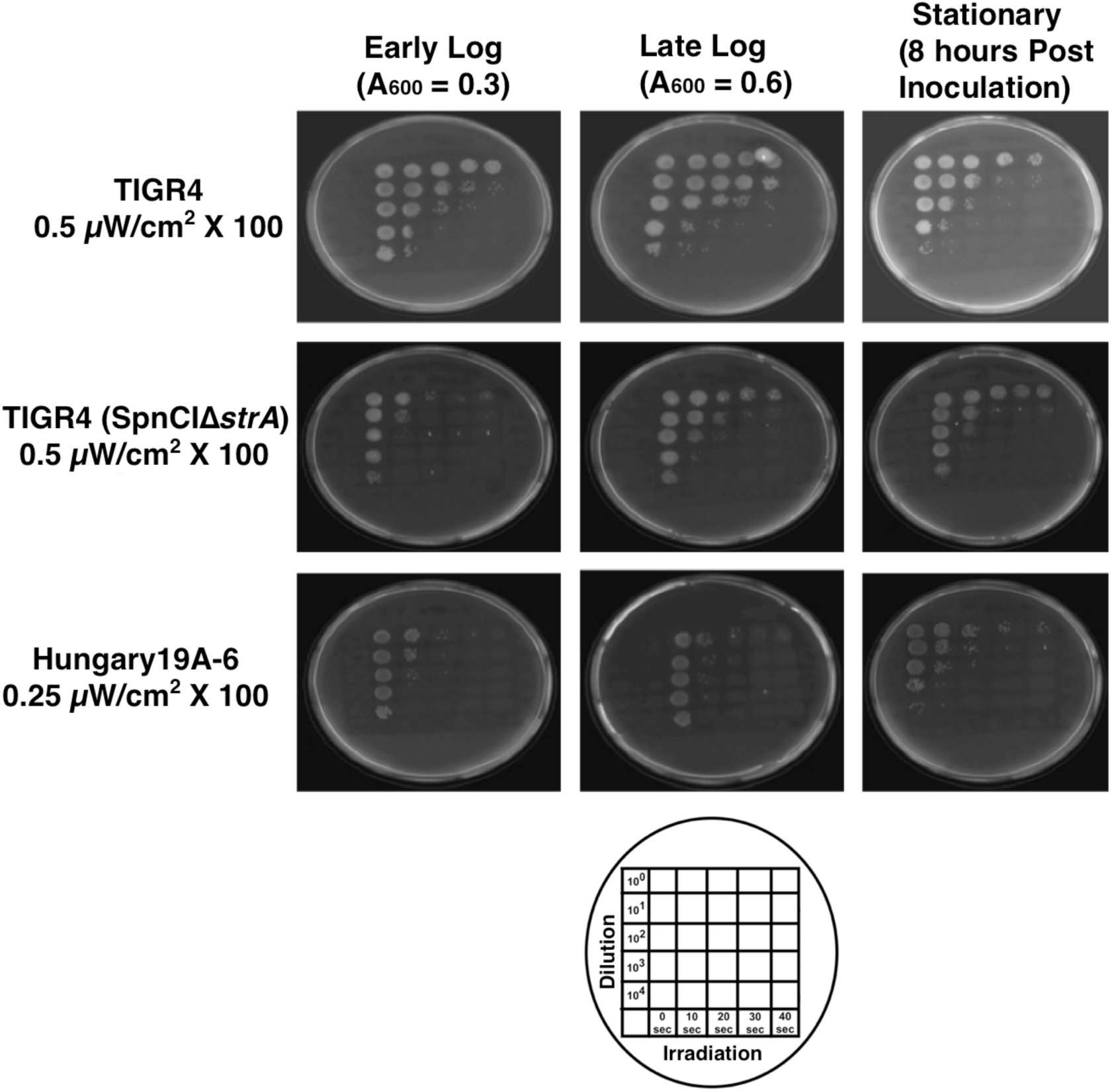
SpnCI integration leads to enhanced UV susceptibility. UV susceptibility of TIGR4 (top row) and isogenic pair TIGR4 SpnCIΔ*strA* (middle row) were compared to Hungary19A-6 (bottom row) at early log (left column), late log (middle column), and stationary (right column). Spots (20μL) were exposed to 0-40 seconds of UV (see plate scheme chart), wrapped to block light and incubated overnight. TIGR4 SpnCIΔ*strA* was more susceptible to UV compared to the wildtype having a more similar susceptible phenotype to SpnCI wildtype containing strain Hungary19A-6.

### Virulence and inflammation studies

For *Galleria mellonella* infections, morbidity and mortality was monitored out to five days post infection with either TIGR4 SpnCIΔ*strA* (red) or TIGR4 (grey) and included a media-only control group (black). Ten 5^th^ instar caterpillars per treatment group were included with three biological repeats (totaling 30 larvae per treatment). Infections with the SpnCI containing strain of TIGR4 were reproducibly greater in both morbidity (p<0.05) and mortality (p<0.05). Specifically, caterpillars infected with the SpnCI containing strain had an overall survival of only 20% and mean health score of 1.6 at day five compared to 50% and 3.85 in the SpnCI null TIGR4 infections (see Figure 5 A&B). To determine whether SpnCI elicited more of an inflammatory response during infection, transcript levels of four antimicrobial peptides (AMP) were measured: gallerimycin (dark grey), galiomycin (red), insect metalloproteinase inhibitor (IMPI; light grey) and lysozyme (green) four hours post-infection. Levels of galiomycin, IMPI, and lysozyme remained similar amongst the two infections. Only gallerimycin was consistently elevated in SpnCI infections, around 4-fold higher, compared to wildtype TIGR4 infections (see Figure 5C). Measurements of bacterial burden in *G. mellonella* at 4, 8, and 24 hours post-infection were collected. TIGR4 burdens were higher at 4 and 24 hours compared to TIGR4 SpnCIΔ*strA* infections. Due to high amounts of variability, no statistical difference between infections were observed (see Figure 5D).

**Figure 5.**
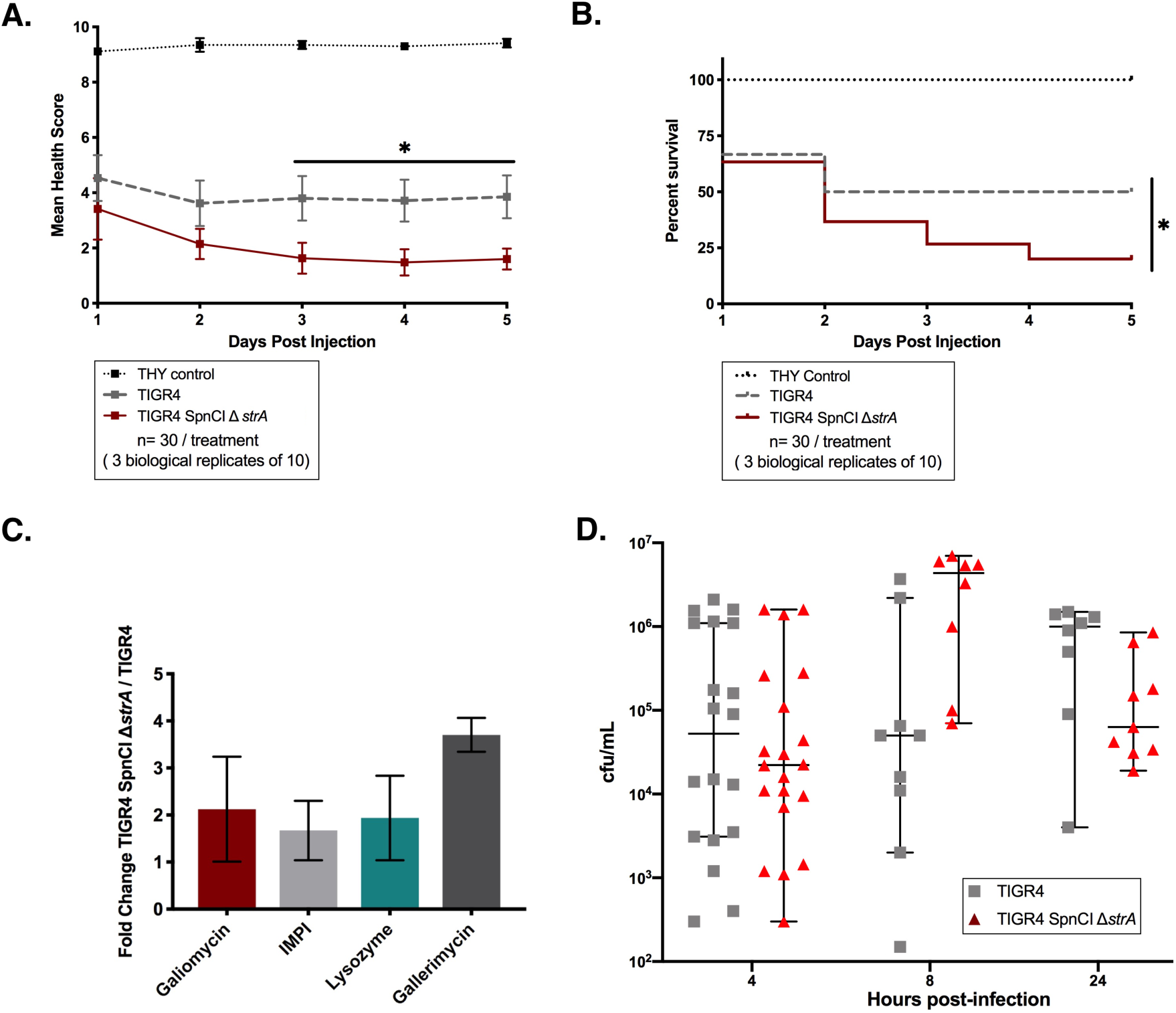
SpnCI enhances virulence in the invertebrate acute infection model *Galleria mellonella.* Fifth instar larvae were injected with ∼10^6^ total cfu of either TIGR4 (grey) or TIGR4 SpnCIΔ*strA* (red). A media-only group was included as a control (black). Data shown is comprised of three biological replicates using ten larvae per group (n=30 total per group). **A.** SpnCI led to significantly higher morbidity measurements by day three compared to TIGR4 (SpnCI null) wildtype (* p < 0.05; 2-way ANOVA with Tukey’s multiple comparisons; means with the standard error of mean shown). **B.** Mortality was significantly higher in SpnCI containing TIGR4 infections compared to the wildtype null strain (* p < 0.05; Mantel-Cox Log Rank). **C.** Four hours post injection, RNA was extracted from larvae and RT-qPCR was performed for four known inflammatory markers: galiomycin (red), insect metalloproteinase inhibitor (IMPI) (light grey), lysozyme (blue), and gallerimycin (dark grey). Only gallerimycin was observed to be consistently higher in SpnCI infected larvae compared to wildtype (dark grey) (mean fold changes plotted with standard error of means). **D.** Measured bacterial burdens of *G. mellonella* at 4, 8, and 24 hours post infection for TIGR4 (grey) and TIGR4 SpnCIΔ*strA* infections (red). Median cfu/mL and 95% CI interval shown.

### SpnCI alters transcriptional patterns

Genes differentially expressed 3-fold or greater with a false discovery rate of less than or equal to 0.05 were considered significant. Triplicate biological repeats of both TIGR4 and TIGR4 SpnCIΔ*strA* were included. SpnCI was observed to mediate changes to transcriptional profiles during late logarithmic growth. In particular, downregulation of capsule and operons involved with secretion and modification of surface adhesins were observed in TIGR4 SpnCIΔ*strA* (see Figure 6). Upregulation of *corA* and shared operon gene SP_0184 in TIGR4 SpnCIΔ*strA* occurred, suggesting SpnCI integration favors *corA* expression. *UvrA* was slightly downregulated in TIGR4 SpnCIΔ*strA* compared to TIGR4 but not significantly enough to be included into the compilation.

**Figure 6.**
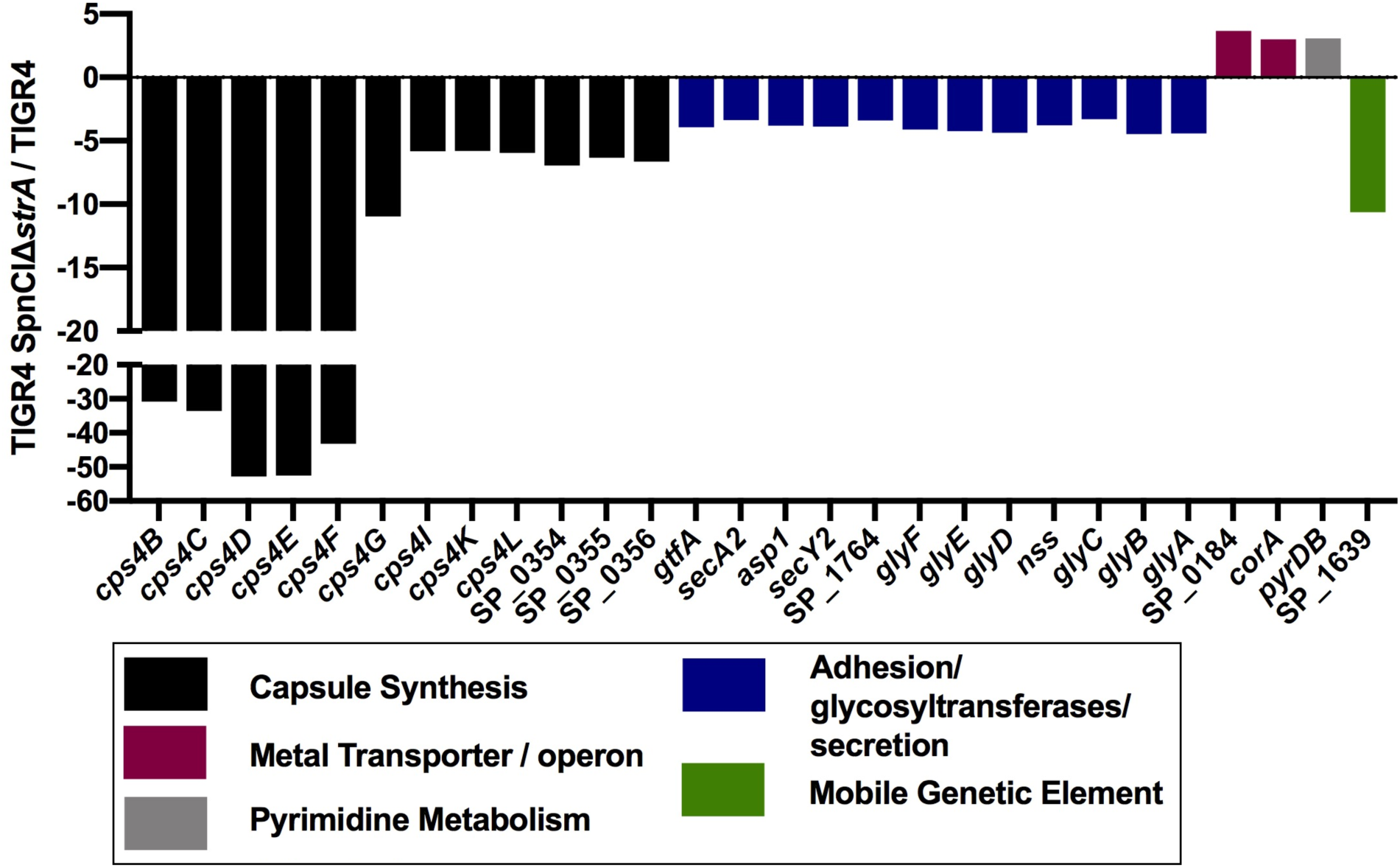
SpnCI alters transcription. RNA sequencing included biological triplicates of TIGR4 and TIGR4 SpnCIΔ*strA*. Genes with fold changes of 3 or greater with a false discovery rate of 0.05 or lower were considered significant. Designated fold change values represent changes observed in TIGR4 SpnCIΔ*strA* in comparison to TIGR4. SpnCI mediated downregulation of operons and genes that are involved with modeling the cell surface. Both capsule (black) and surface modeling associated (blue) operons were down regulated. *CorA* and the accessory gene SP_0184 (purple) were up suggesting SpnCI integration favors *corA* expression.

### Resident prophages of Hungary19A-6 and dissemination of SpnCI

SpnCI was detected from mitomycin C induced and purified Hungary19A-6 Δ*strA* phage lysate (see Figure 7A) but not from TIGR4 SpnCIΔ*strA* (see Figure 7B). As Hungary19A-6 has two resident prophages but TIGR4 does not, these results suggest that SpnCI uses a helper prophage for packaging and dissemination. Both annotated Hungary19A-6 prophages were also detected within the lysate. Electron microscopy of Hungary19A-6 Δ*strA* revealed an average capsid size of 60 nm of typical siphoviridae morphology (see Figure 8A). Two of the sixty measured had a substantially smaller capsid size (see Figure 8B). Electron microscopy of the TIGR4 SpnCIΔ*strA* did not reveal any phage particles, confirming the negative results observed by PCR screen and lack of annotated prophages within the TIGR4 wild type genome (GenBank accession AE005672.3). To determine whether the packaged SpnCI particles were infectious, TIGR4 and capsule null mutant T4R, were co-incubated with the SpnCIΔ*strA* containing lysate at either early or late logarithmic growth for 15- or 60-minutes. SpnCIΔ*strA* infection experiments revealed successful infections only within the T4R strain both at 15- and 60-minutes co-incubation (see Table 2). TIGR4 under the same conditions failed to yield SpnCI infections. To rule out transformation as cause of horizontal gene transfer, inactivated phage (boiled lysate) was co-incubated for an hour with either TIGR4 or T4R strains, plated and screened for SpnCI Δ*strA*::*kanR* containing strains. No kanamycin resistant colonies were observed, ruling out transformation.

**Table 1.** Strains of *Streptococcus pneumoniae* used in this work.

| Strain | Characteristics | Reference |
| --- | --- | --- |
| Hungary19A-6 | Serotype19A multidrug resistant clinical genome isolate | (8) |
| Hungary19A-6Δ*strA* | Isogenic pair of Hungary19A-6 with *kanR* tagged phage-like element SpnCI | This work |
| TIGR4 | Serotype 4 clinical genome isolate | (45) |
| T4R | Capsule null strain due to polar mutation of capsule operon via insertion of otherwise identical TIGR4 genetic background | (37, 38) |
| T4R SpnCIΔ*strA* | T4R strain with *kanR* tagged phage-like element SpnCI | This work |
| TIGR4 SpnCIΔ*strA* | Capsule rescued strain with *kanR* tagged phage-like element SpnCI | This work |

**Table 2.**
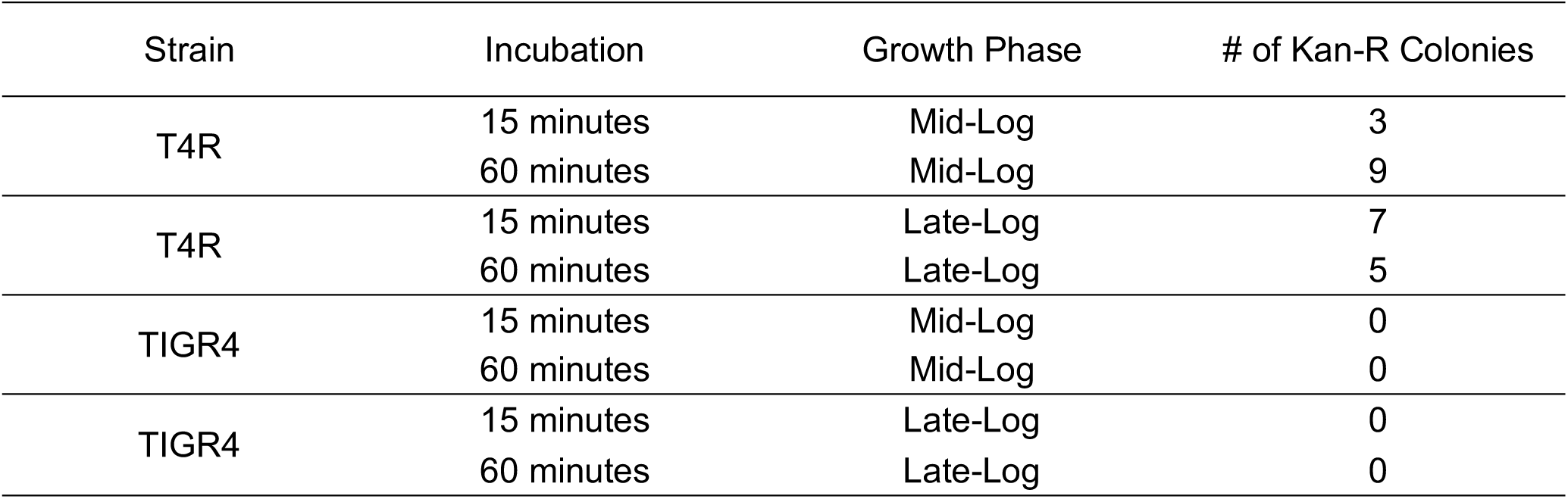
SpnCI infection inhibited by capsule

| Strain | Incubation | Growth Phase | # of Kan-R Colonies |
| --- | --- | --- | --- |
| T4R | 15 minutes | Mid-Log | 3 |
|  | 60 minutes | Mid-Log | 9 |
| T4R | 15 minutes | Late-Log | 7 |
|  | 60 minutes | Late-Log | 5 |
| TIGR4 | 15 minutes | Mid-Log | 0 |
|  | 60 minutes | Mid-Log | 0 |
| TIGR4 | 15 minutes | Late-Log | 0 |
|  | 60 minutes | Late-Log | 0 |

**Figure 7.**
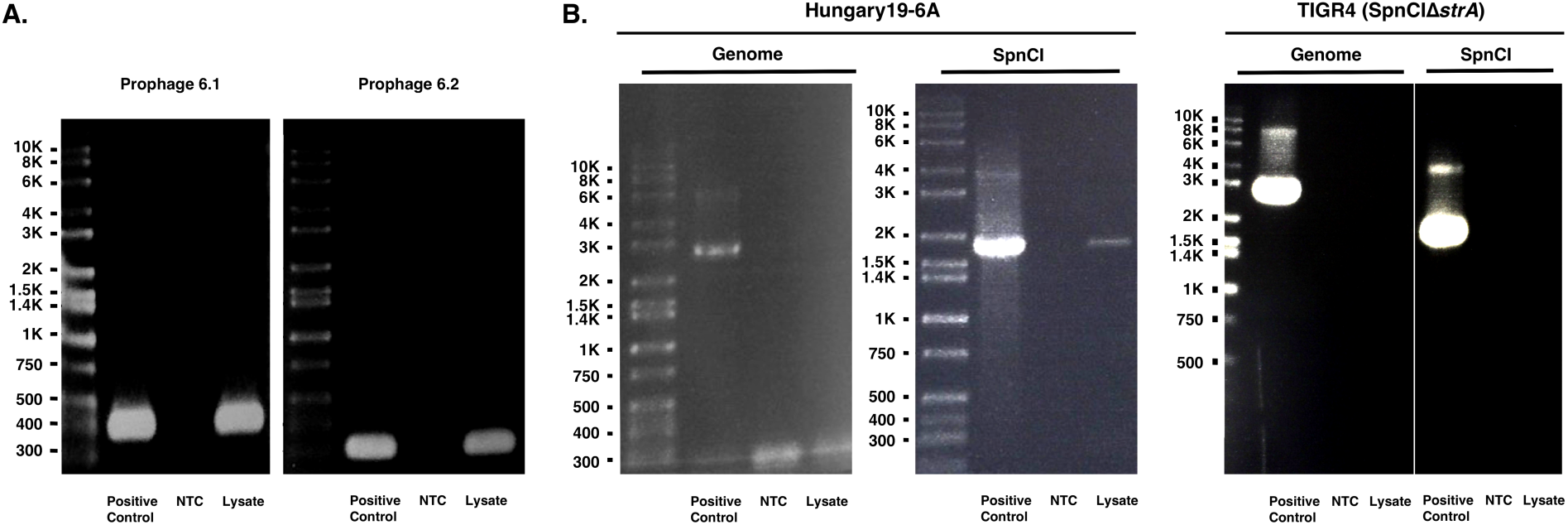
SpnCI is detected in phage lysate when resident prophages are present. Mid-log cultures were treated with 0.5 mg/mL of mitomycin C and left overnight before isolating phage by centrifugation. Lysate was treated with DNase to remove exogenous DNA then boiled to release phage DNA. **A.** Both annotated resident prophages 6.1 and 6.2 were detected in phage lysate from Hungary19A-6. An unexpected find was the presence of SpnCI DNA, suggesting that SpnCI can utilize resident phage structural genes in order to package itself. No contaminant bacterial DNA was detected. **B.** TIGR4 has no annotated resident prophages. As such, no SpnCI was detected in TIGR4 SpnCIΔ*strA* mitomycin C treated samples.

**Figure 8.**
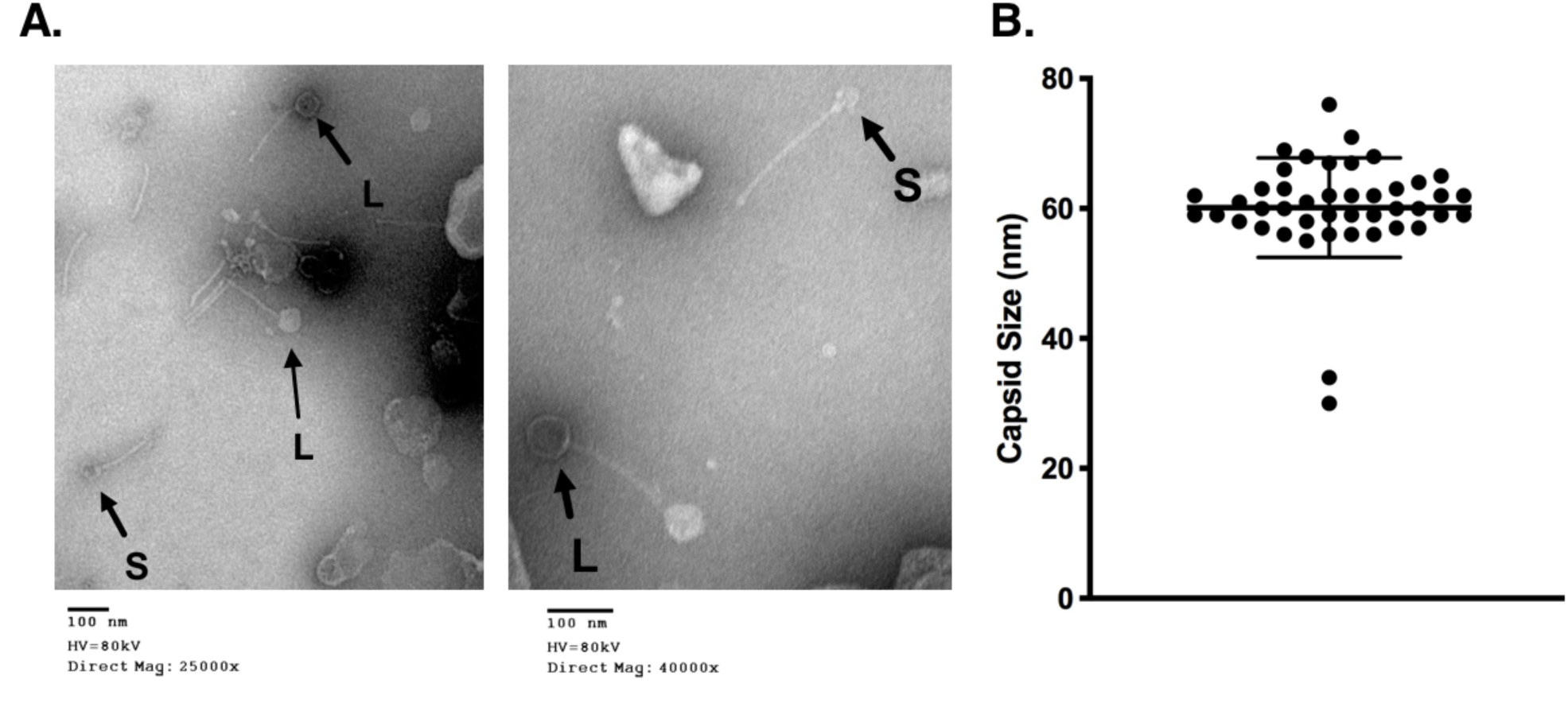
Smaller capsid heads are observed suggesting restructuring of capsid for SpnCI packaging. **A.** Electron microscopy of Hungary19A-6 Δ*strA* revealed phage with typical siphoviridae morphology. A smaller (S) and larger (L) size capsid was observed with the larger forms being more prevalent. **B.** Capsid size averaged around 60 nm however there were two with much smaller sizes ranging around 35-37 nm in size.

## Discussion

Our data suggests that under laboratory conditions SpnCI remains predominantly integrated and does not undergo growth dependent excision and integration patterns as previously described in *S. pyogenes* (5). This integrated state was found to have the phenotypic effect of enhanced UV sensitivity in TIGR4 SpnCIΔ*strA* compared to the SpnCI null wildtype strain TIGR4, mirroring the UV susceptibility patterns observed in Hungary19A-6. Further *in vivo* studies are necessary in order to capture the biological generation of excision / integration flux as it remains unlikely from a cost fitness perspective for strains containing SpnCI to be continuously deficient in nucleotide excision repair. We performed the integration excision studies using Hungary19A-6 (SpnCI wildtype strain) in order to account for the potential influence other resident prophages may have on SpnCI integration and excision patterns.

Although bacterial burdens were not significantly different amongst TIGR4 and TIGR4 SpnCIΔ*strA* infections, SpnCI infections caused significantly greater morbidity and mortality measurements within the invertebrate acute infection model *Galleria mellonella*. This phenomenon was most evident at 3 days and sustained until the end time point of 5 days post infection. Bacterial burdens were assessed out to 24 hours post infection due to the high mortality that occurs after day 1. Bacterial burdens may differ amongst the two strains 3 days and onward but the low sample numbers and high amount of variability of cfu/mL recovered makes assessment of burden challenging. Although there was a marked increase in morbidity and mortality with SpnCI infections, inflammation transcript levels for three of the four measured inflammatory markers were highly variable when compared to the TIGR4 (SpnCI null) infected larvae. Interestingly, transcript levels only for gallerimycin were consistently raised, around 4-fold higher in SpnCI infected larvae. Gallerimycin is known for its anti-fungal properties but has also previously been described as inducible in the presence of lipopolysaccharide (lps) [205]. Recombinant gallerimycin was found to only have inhibitory activity against entomopathogenic fungus *Metarhizium anisoliae* while no effect was observed with either Gram-positive, Gram-negative or yeast (24). As authors have previously noted, the killing activity of gallerimycin was only assessed using a recombinant peptide and to our knowledge the question of whether post translational modification of this peptide influences activity against other non-fungal organisms remains unknown. It is interesting to note that gallerimycin was consistently up-regulated in larvae infected with the SpnCI containing strain of TIGR4 as compared to the wild type SpnCI null TIGR4 infections. In an attempt to elucidate potential immunological signaling pathways, we searched for transcription factor binding sites that were unique to gallerimycin utilizing previously described conserved NF-κB, C/EBP and CRE-BP1 sequences of other lepidopteran species (24, 25). A putative NF-κB binding site was only identified in gallerimycin, suggesting that SpnCI may be inducing an immunological pathway involving NF-κB.

RNA sequencing during late logarithmic growth demonstrated SpnCI’s involvement with downregulation of multiple genes that are involved with cell surface modelling. This included marked downregulation of the capsule operon (see black in Figure 6). Capsule shedding has been reported to occur during invasive infection through enabling respiratory epithelial adhesion, a precursor to invasive disease (26). A more modest downregulation of surface modelling operons that are involved with glycosylation and surface transport of pneumococcal serine-rich repeat adhesins (see blue in Figure 6) (PsrP) (27, 28) was observed. These included the accessory sec system (secA2 and secY2) which is involved in adhesion as well as the export of pneumolysin toxin and biofilm formation (29). It is interesting that the glycosyltransferases and accessory sec system are down in TIGR4 SpnCIΔ*strA* compared to TIGR4 as these genes have been shown to enhance virulence (28) yet in our invertebrate infection model, TIGR4 SpnCIΔ*strA* was significantly more virulent. Our observations suggest that downregulation of capsule is sufficient for enhancing virulence, at least within the invertebrate infection model. Further virulence studies using the murine model are necessary to assess the contribution of SpnCI to enhanced virulence. Our studies are the first to demonstrate the phenomenon of transformation-independent movement of phage-like elements within streptococci, suggesting that the occurrence of prophage hijacking extends beyond what has previously been described in *Staphylococcus aureus* (30, 31). EM studies demonstrated only 2 of the total 60 measured capsids were markedly smaller in size, averaging around 32 nm, as compared to the average 60 nm. Further work is necessary to determine if lower frequency of SpnCI is due to packaging efficiency of SpnCI or simply due to the fact that two other prophages are also present, reducing the observed frequency of smaller sized heads. The initial susceptibility studies demonstrate that capsule impedes SpnCI infection as no SpnCI infected colonies were observed in TIGR4, but were observed in capsule mutant strain T4R (see Table 2). In other recent studies, the process of capsule shedding has been described and indicates that during invasion of epithelial cells, pneumococcus uniformly sheds capsule through LytA activity (26). During the phenomenon of capsule shedding, cells potentially become more susceptible to SpnCI infection, mediating further spread of SpnCI in otherwise seemingly resistant encapsulated strains.

Overall, our research has demonstrated the implications of carrying pneumococcal phage-like element. When present SpnCI: enhanced virulence, altered UV susceptibility, modulated transcriptional patterns, and was demonstrated capable of dissemination via infection of susceptible pneumococci. Further research is necessary to identify distinct environmental mediators of SpnCI excision within an *in vivo* setting and to determine what SpnCI specific gene(s) enhance pneumococcal virulence. We also plan to utilize a murine infection model to investigate the potential role SpnCI may play in modulating the host immune response during infection. Ultimately, a better understanding of SpnCI’s role in virulence may lead to the development of diagnostics and or therapeutics for severe pneumococcal infections.

## Acknowledgements

We are grateful to Dr. Justin Thornton from Mississippi State University for donating null capsule mutant T4R and strain DP1617. We also would like to acknowledge Dr. Thornton, Dr. Don Morrison, and Dr. Roger Junges for help with optimization of the transformation protocols. We thank Dr. Allison Gillaspy and Ms. Jenny Gipson from the Laboratory for Molecular Biology and Cytometry Research at OUHSC for assistance in the sequencing of the generated T4R SpnCIΔ*strA* genome, other confirmation sequencing for capsule rescue, and for the RNA sequencing experiment. We also thank Mr. Ben Fowler and the Oklahoma Medical Research Foundation Imaging Core Facility, for providing bacteriophage staining and electron microscopy services. We are also grateful to Dr. Hariprasad Gali for synthesis of the CSP-1 and CSP-2 peptides. Research reported in this publication was supported by the National Institute of General Medical Sciences of the National Institutes of Health under award number GM103648.

## Materials and Methods

### UV SpnCI induction

The frozen stock of S. pneumoniae Hungary19-6A (0.5 ml) was diluted into 25 ml THY supplemented with 2.5 ml lysed horse blood. The culture was incubated at 37° C until A600 nm > 1.0. Two 10 ml were dispensed into sterile Petri dishes, and one dish was irradiated at UV 260 nm for 45 seconds in a darkened room and then wrapped in foil to prevent photoreactivation. The other culture served as an untreated control. Both dishes were then incubated at 37°C for 30 minutes. The cells were harvested by centrifugation, resuspended in 0.5 ml RNAlater, and stored at 4°C overnight. The next day, DNA was harvested from each cell pellet as previously described (4, 5, 32). PCR was then performed using the primers described in Supplemental Table 1 on each DNA sample to detect the presence of *attP* and *attB*, indicative of the excised SpnCI, or *attL* and *attR*, which appear following SpnCI integration into the bacterial chromosome.

### Generation of isogenic pairs

Wild type SpnCI containing strain Hungary19A-6 was transformed with 2,483 bp sized product containing kanamycin resistance (*kanR*) and flanking regions of the *strA* gene generated through PCR fusion (see Figure 1 and Table 3) in order to confer homologous recombination and subsequent knockout of *strA* and tagging with kanamycin resistance [500 μg/mL] (33, 34). Transformation with PCR product was performed in C+Y_YB_ media using a published transformation protocol (35) with CSP-2 (36). To overcome challenges faced while attempting to transform TIGR4 (wild type SpnCI^−^ strain) with the tagged SpnCI *strA*::*kanR* ∼12 kb product, we initially transformed the tagged SpnCI into the capsule mutant T4R (37, 38) (strain generously provided by Dr. Justin Thornton). Transformation was performed using previously described methods with CSP-1 (39, 40). Successfully transformed T4R SpnCIΔ*strA* was sequenced to ensure no other Hungary19A-6 genes were transferred. Capsule rescue was performed by transforming with cps4A amplified product from wildtype TIGR4. The cps4A locus of capsule rescue strain TIGR4 SpnCIΔ*strA* was sequenced in order to confirm capsule rescue in addition to capsule agglutination screening using ImmuLex™ SSI^®^ Pneumococcus Type 4 (Statens Serum Institut, Herredsvejen, Denmark). See table 1 for summary of isolates used.

### SpnCI integration experiments

DNA was collected from Hungary19A-6 every 15 minutes from early log growth (O.D._A600_= 0.3) to stationary phase (O.D._A600_= 0.8) as previously described (5, 32). To remove RNA, samples were treated with 1μL of [10 mg/mL] RNase A before being diluted to a final concentration of [10ng/μL]. 50 ng of DNA was added to the PerfecCTa® SYBR® Green SuperMix (Quantabio, Beverly, MA, USA) master mix. To assess SpnCI integration, qPCR was performed using the Biorad iCycler (BioRad Laboratories, Hercules, CA, USA) (see Table 1 section C for primers and cycling parameter used). To assess abundance, calculations using the ΔΔC_T_ methods were performed using *gyrA* for normalization.

### UV Sensitivity Assay

UV sensitivity of TIGR4, TIGR4 SpnCIΔ*strA*, and Hungary19A-6 were assessed during early log (O.D._A600_= 0.3), late log (O.D._A600_= 0.6), and stationary phase (O.D._A600_= 0.8). Cells were harvested from cultures grown at 37°C with 5% CO_2_ in Todd Hewitt Broth supplemented with 5% yeast extract (THY) at corresponding growth points. Specifically, cells were pelleted by 5,000 rpm centrifugation (at 4°C) for 15 minutes and resuspended in equal volume of sterile 0.1 M MgSO_4_ to maintain initial absorbance. For each strain and time point, a ten-fold serial dilution using sterile 0.1M MgSO_4_ was performed. Onto a Tryptic Soy Agar (TSA) plate supplemented with 5% sheep blood, 2 μL drops of undiluted and serially diluted cultures were added (see Figure 4A for plating scheme). Drops were then exposed to UV for a duration of time ranging from 10 to 40 seconds. An unexposed row was included as a growth control. For both TIGR4 and TIGR4 SpnCIΔ*strA* a UV dosage of 0.25 μW/cm^2^ x100 was used. Hungary19A-6 was inherently more UV sensitive, a lower exposure of 0.25 μW/cm^2^ x100 was necessary in order to not completely inhibit growth. After exposure, plates were immediately wrapped in foil (to prevent photoreactivation of DNA repair) and incubated overnight using previously described growth conditions. Photos were taken the following day using the Flash and Go Automatic Colony Counter (IUL Instruments, Barcelona, Spain).

### Virulence and inflammation studies

TIGR4 and TIGR4 SpnCIΔ*strA* rescue strains were grown in Todd Hewitt Broth supplemented with 5% yeast extract (THY) at 37°C with 5% CO_2._ Once cultures reached O.D._A600_ = 0.6, cultures were spun for 15 minutes at 5,000xg, 4°C and resuspended to an O.D._A600_ = 0.3. Fifth instar *Galleria mellonella* weighing 0.3 to 0.6 grams were chilled at 4°C for ten minutes prior to receiving the 5 μl injection within the hindmost right pro-leg (6.25×10^5^ to 1.2×10^6^ cfu total). Three biological replicates with ten larvae per group were performed for each treatment group (total of 30 caterpillars per treatment). A THY only inoculum group was included as a control. Larvae were monitored post injection for five days and assessed each day for mean health and survival using previously described criterion (41, 42). To ensure the intended inoculum concentration was achieved, concentrations were quantified using the simplified agar plate method (43). For inflammation studies, infections were performed as described above. After 4 hours post infection, waxworms were anesthetized at 4°C for ten minutes then ground in 1 mL of TRIzol™ (Invitrogen™, Carlsbad, CA) using the sterile 15 mL conical tissue grinder (VWR™, Radnor, PA) and RNA extraction was performed according to manufacturer’s protocol. For qPCR, gallerimycin, galiomycin, lysozyme and insect metalloprotease transcript levels were measured using previously reported methods (44) with SYBR green dye and 10 ng of cDNA. Experiments consisted of 10 caterpillars per strain with 3 biological repeats (30 caterpillars per infection). Bacterial burdens of TIGR4 and TIGR4 SpnCIΔ*strA* were measured at 4, 8, and 24 hours post infection. Infected larvae were anesthetized at 4°C for ten minutes, ground with 1 mL of phosphate buffered saline using the sterile 15 mL conical tissue grinder (VWR™, Radnor, PA). Samples were spun at 5,000xg, 4°C for ten minutes to sediment debris and serial dilutions of supernatant were plated onto [50μg/mL] nalidixic acid containing Tryptic Soy Agar (TSA) plate supplemented with 5% sheep blood. Plates were incubated at 37°C with 5% CO_2._ overnight prior to measuring bacterial burdens.

### RNA Sequencing

RNA was isolated (growth conditions described above), from late logarithmic cultures (O.D._A600_= 0.6). Total RNA extraction was performed using the Direct-zol™ RNA MiniPrep Plus (Zymo Research, Irvine, CA). The Ribo-Zero rRNA Removal Kit (Bacteria) (illumina^®^, San Diego, CA) was used for mRNA enrichment and the RNA Clean & Concentrator™ −5 (Zymo Research, Irvine, CA) for final purification. Quality of enriched mRNA samples was confirmed using the Agilent 2100 BioAnalyzer (Agilent Technologies, Inc., Santa Clara, CA, U.S.A.) RNAseq libraries were constructed using the Illumina TruSeq RNA LT v2 kit and established protocols. The library construction was done using ribosomally depleted *Streptococcus pneumoniae* RNA (500ng). Each library was indexed during library construction in order to multiplex for sequencing on the Illumina MiSeq platform. Samples were normalized and the libraries were pooled onto 2, 2×150bp paired end sequencing runs on the Illumina MiSeq. A total of 55 million reads of sequencing data was collected for the samples. Raw data for each sample was analyzed using CLC Genomics Workbench version 11.0.1 software from Qiagen (formerly CLCBio). Raw sequence reads were mapped to the TIGR4 *Streptococcus pneumoniae* genome (Accession AE005672.3) for identification of genes expressed under each condition. Pairwise comparison of the expression results were performed using the total mapping results for TIGR4 SpnCIΔ*strA* vs TIGR4. Differential gene lists were created with 3-fold expression cutoff and significant p values of less than or equal to 0.05 with a false discovery rate of 0.05 or less to identify genes that were up- or down-regulated under each condition. RNA sequencing data was compiled from triplicate repeats. The transcriptional analysis data is available through accession GSE132556.

### Prophage and SpnCI induction and electron Microscopy

Strains Hungary19A-6 and TIGR4SpnCIΔ*strA* were grown as described above until mid-log growth O.D. _A600_= 0.6. Cultures were induced using a concentration of 0.5 mg/mL of mitomycin C and incubated overnight at 37°C with 5% CO_2_. An initial centrifugation was performed to pellet bacteria (5,000 rpm for 15 minutes a 4°C) then supernatant was filtered (0.45 μm). The resulting lysate was further spun (40,000xg for 2 hours at 4°C) and pellet resuspended in THY broth. Residual DNA was removed by [7.5 μg/mL] DNase I treatment (Sigma-Aldrich, St. Louis, MO.) for 30 minutes at 37°C. Lysate was screened for contaminant bacterial DNA by PCR. Electron microscopy was performed at Oklahoma Medical Research Foundation’s Imaging Core Facility (Oklahoma City, Oklahoma) using the Hitachi H-7600 Transmission Electron Microscope. Phage particles were stained using the following protocol: 10 μL of sample was settled either for 1, 2 or 4 minute(s) on a 300 hex-mesh, formvar coated, glow discharged copper grid using the single drop method with excess being wicked with filter paper. The grid was washed with DI H_2_O for 10 seconds, excess water removed and then stained with 10 μL of 5% ammonium molybdate 1% trehalose in DI H_2_O (pH 7.01) for 1 minute. Staining solution was removed washed and allowed to air dry for 1 minute.

### SpnCI infection experiments

TIGR4 and T4R were grown at 37°C with 5% CO_2_. A 1:1 ratio of lysate obtained from Hungary19A-6 Δ*strA*::*kanR* (method described above) to TIGR4 or T4R cultures was incubated for either 15 minutes or 1 hour at mid-log or late-log stage of bacterial growth at 37°C in the presence of 5% CO_2_. The mixture was then plated onto a [500 μg/mL] kanamycin Tryptic Soy Agar (TSA) plate supplemented with 5% sheep blood. Colonies were counted after overnight incubation and screened by PCR to confirm the presence of SpnCIΔ*strA* (see Table 1 section C). As a control for transformation, co-incubation with inactivated (boiled) lysate was performed; no kanamycin resistant colonies were observed.

---

**Supplemental Table 1.**
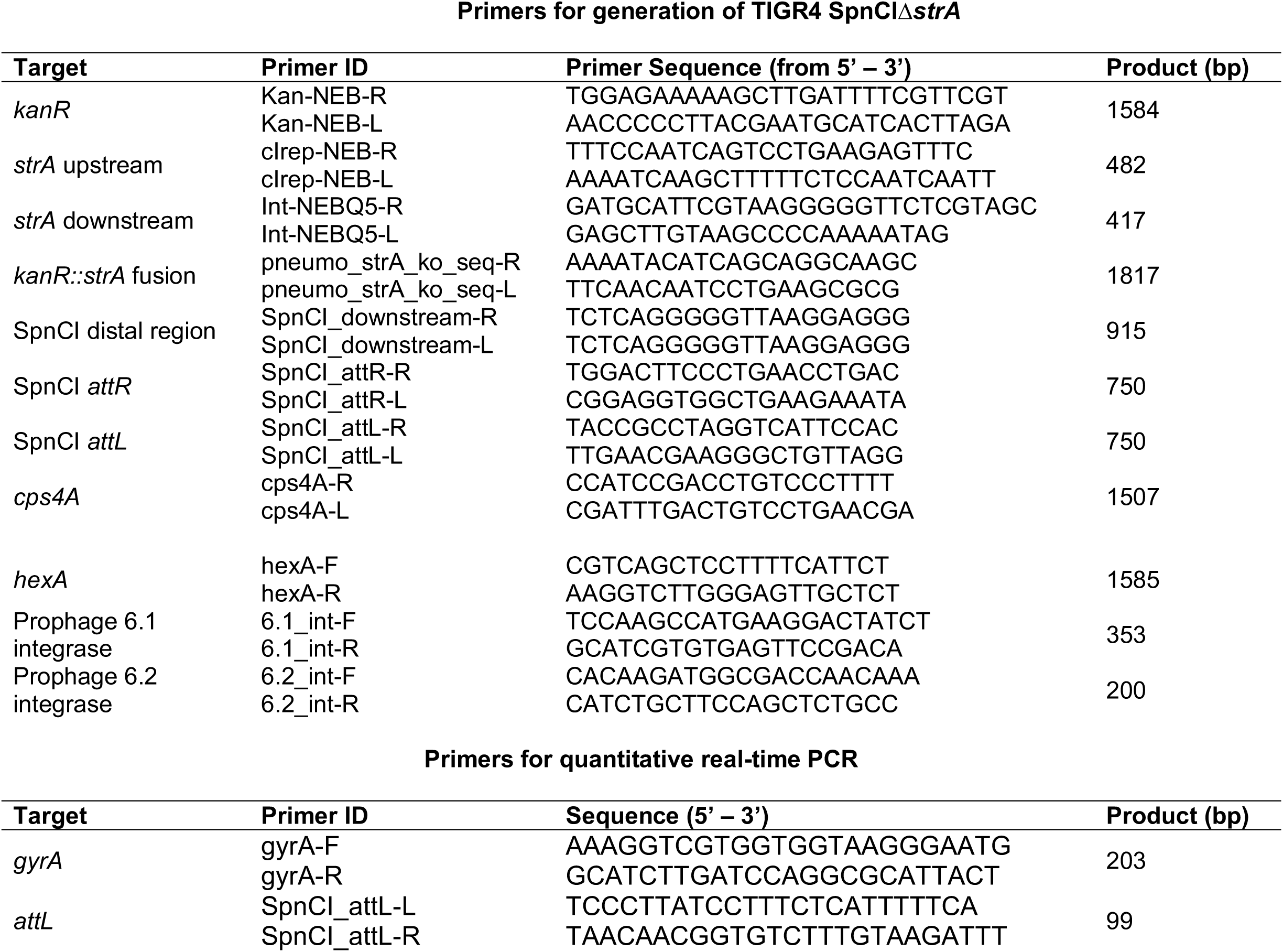
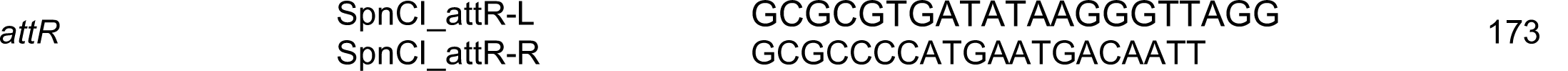
PCR primers used in this work

